# High-coverage allele-resolved single-cell DNA methylation profiling by scDEEP-mC reveals cell lineage, X-inactivation state, and replication dynamics

**DOI:** 10.1101/2024.10.01.616139

**Authors:** Nathan J. Spix, Walid Abi Habib, Zhouwei Zhang, Emily Eugster, Hsiao-yun Milliron, David Sokol, KwangHo Lee, Paula Nolte, Jamie Endicott, Kelly F. Krzyzanowski, Toshinori Hinoue, Jacob Morrison, Benjamin K. Johnson, Wanding Zhou, Hui Shen, Peter W. Laird

**Author notes:** Equal contribution (Nathan J. Spix, Walid Abi Habib).

## Abstract

DNA methylation is a relatively stable epigenetic mark with important roles in development and disease.^1^ Since cell-to-cell variation in epigenetic programming can reflect important differences in cell state and fate, it is clear that single-cell methods are essential to understanding this key epigenetic mark in heterogeneous tissues. Existing single-cell whole-genome bisulfite sequencing (scWGBS) methods^2^ have significant shortcomings, including very low CpG coverage^3–7^ or inefficient library generation requiring extremely deep sequencing.^8^ These methods offer limited insight into focal regulatory regions and generally preclude direct cell-to-cell comparisons. To address these shortcomings, we have developed an improved method, scDEEP-mC (single-cell Deep and Efficient Epigenomic Profiling of methyl-C). We show that high-coverage promoter methylation profiling by scDEEP-mC can identify multiple cell types, while allele-resolved methylation calls allow assessment of X-inactivation state in single cells and identification of transcription factor binding sites (TFBS) enriched for hemi-methylation. We also use scDEEP-mC to profile single-cell copy number alterations, identify actively replicating cells, and track DNA methylation dynamics during and after DNA replication. The high coverage of scDEEP-mC creates an exceptional opportunity to explore DNA methylation biology in individual cells.

## Main Text

scDEEP-mC is optimized to provide high coverage at moderate sequencing depth by efficient production of complex libraries. This facilitates direct cell-to-cell comparisons and provides insights into biology that is obscured by the cluster-wise comparisons or megabase-scale summarization necessitated by low-coverage methods. Our method is based on the post-bisulfite adapter tagging approach,^9^ which usually incorporates a cleanup step after bisulfite conversion to replace high-concentration NaHSO_3_ with a buffer less hostile to polymerase activity. However, this can result in significant DNA loss, especially with low input DNA quantities. To prevent such loss, we sort cells directly into a small volume of high-concentration NaHSO_3_. Following bisulfite conversion, this solution is diluted to a concentration low enough to allow polymerase activity. We also improved on existing methods by designing our random primers to complement the expected base composition of the primed fragments and carefully titrating the concentration of these primers to mitigate adapter contamination and other technical artifacts (Fig. 1a).

**Figure 1.**
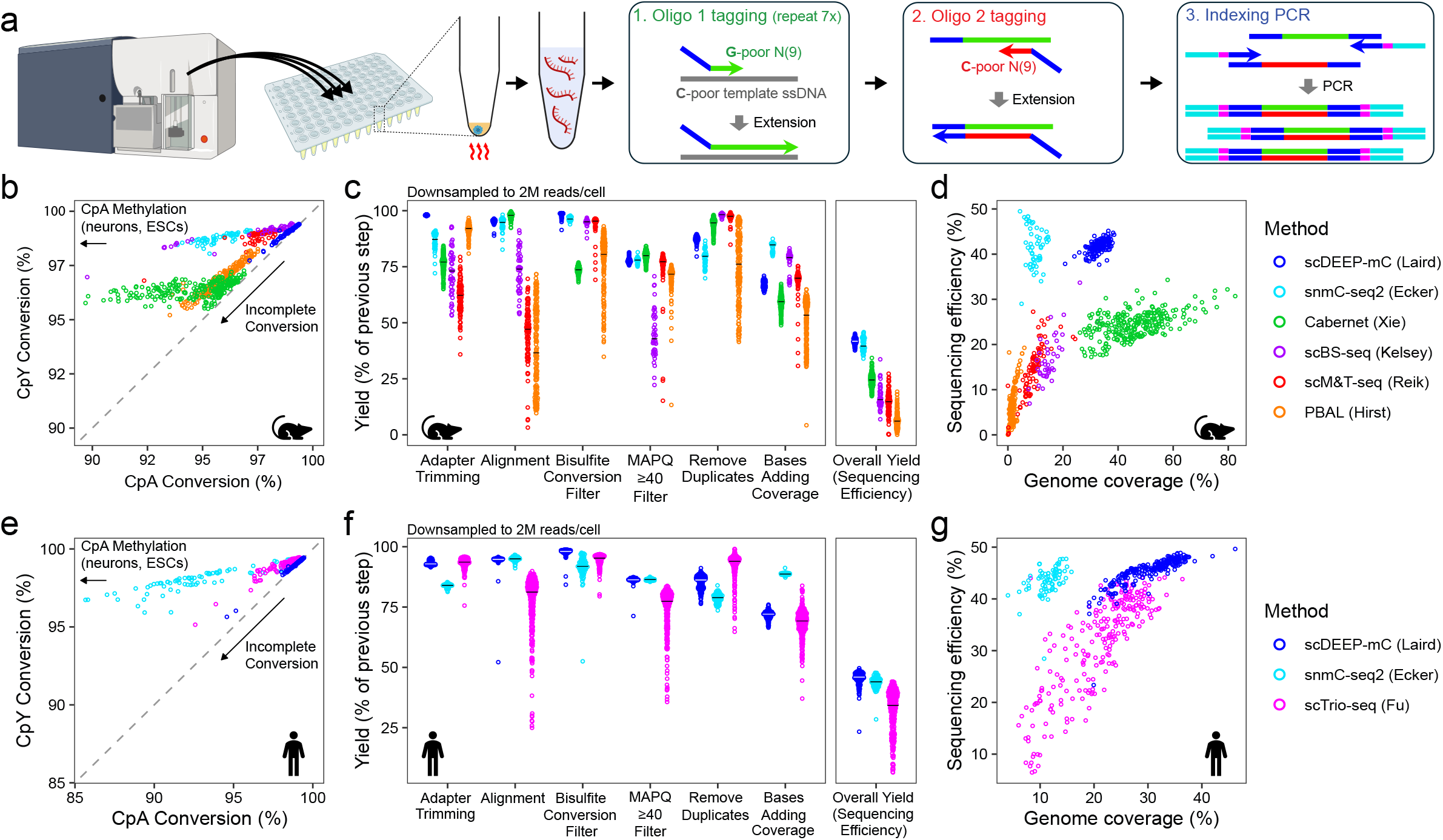
scDEEP-mC provides high coverage in single cells by efficient production of high-complexity libraries. (a) Library preparation overview. Single cells derived from culture or primary tissue are flow-sorted into bisulfite conversion buffer in a 96-well plate. Heating facilitates cell lysis, bisulfite conversion, and DNA fragmentation in one step. A 20x dilution step replaces purification after bisulfite conversion, eliminating DNA loss. First-strand adapters are added via seven rounds of base-composition-matched random priming, followed by second strand adapter tagging (also via random priming) and indexing PCR. (b-g) Comparison of scDEEP-mC with other scWGBS protocols. Publicly available raw sequence data from mouse (b-d) and human (e-g) were downloaded from^2–8^ and analyzed using a standardized, Biscuit-based^10^ pipeline (see Methods). Potential doublets and hyperdiploid cells were excluded. (b, e) scDEEP-mC displays reliable, complete bisulfite conversion, as evidenced by consistently high CpY conversion. (c, f) 2 million reads were randomly sampled from each cell and the number of bases remaining after each processing step was recorded. From left to right, processing steps include removal of sequencing adapters, mapping, removal of reads with ≥ 2 CpY retention events, removal of reads with MAPQ < 40, and removal of duplicate reads. ‘Bases adding coverage’ denotes the proportion of reads that do not overlap with other reads in the final dataset, and ‘overall yield’ represents the number of covered bases per sequenced base. The median of each distribution is noted with a crossbar. scDEEP-mC displays extremely low adapter contamination and high alignment rates (especially compared to scBS-seq, to which it is most similar). Overall, scDEEP-mC achieves the highest efficiency of all compared methods. (d, g) Overall sequencing efficiency (from panels c and f) and genomic coverage in all compared cells. scDEEP-mC has the highest efficiency of any high-coverage method.

We performed scDEEP-mC on 175 primary cells isolated from mouse intestinal epithelium and 292 cultured primary human fibroblasts and evaluated its performance in comparison to publicly available scWGBS datasets. Of the many methods currently available, we focused on those with the highest reported CpG coverage. To avoid biases introduced by widely varying analysis methodologies, we processed all raw sequencing datasets through a standardized analytical pipeline based on Biscuit^10^. We evaluated the bisulfite conversion rate (Fig. 1b, e), sequencing efficiency (bases covered per base sequenced [Fig. 1c, f]), and genomic coverage (Fig. 1d, g) of these methods, excluding hyperdiploid cells and doublets (see Methods). scDEEP-mC displays consistently high bisulfite conversion rates (Fig. 1b, e), minimal adapter contamination, and very high alignment rates (Fig. 1c, f). Overall, our results show that at a sampling depth of 2M reads/cell, about 40% of bases sequenced by scDEEP-mC contribute unique coverage, indicating that scDEEP-mC libraries have significantly higher complexity compared to other scWGBS methods. While snmC-seq2^3^ produces libraries of comparable sequencing efficiency (Fig. 1c, f), its very low library yield necessitates pooling many libraries and limits coverage (Fig. 1d, g). The recently developed Cabernet method^8^ achieves high coverage (Fig. 1d), but suffers from high CpY retention due to incomplete cytosine conversion (Fig. 1b) and requires extremely deep sequencing (≥200M reads per cell). By contrast, scDEEP-mC libraries have high yield (facilitating coverage of 30-40% of CpGs, even in primary cells), reach saturation at moderate sequencing depths (35M reads/cell), and display consistently high bisulfite conversion rates.

We explored whether scDEEP-mC could reveal epigenetic information reflecting the distinct biology of individual cells that would be difficult to obtain with lower coverage methods. First, we sought to identify cell types in an unselected population of 175 single primary cells from mouse intestinal epithelium (Fig. 2). We demonstrate three orthogonal methods for identifying cell types: analysis of cell-type-specific hypomethylated regions^11^ (Fig. 2a), unsupervised clustering of variably methylated promoters (Fig. 2b), and dimension reduction using non-negative matrix factorization (NMF) (Fig. 2f). Analysis of cell-type-specific hypomethylated regions is straightforward, but is limited to the broad categories reported in atlases^11^. In contrast, clustering of variably methylated promoters facilitates direct comparison of epigenetic regulatory changes between clusters (Fig. 2c-e). Unsupervised clustering based on promoter methylation levels readily identified two major groups of cells (enterocytes and immune cells [Fig. 2c]), each with two major sub-groups. In one enterocyte subgroup, promoters of genes involved in absorptive processes (such as *Abcg5*) were unmethylated, while in the other subgroup, promoters of genes implicated in differentiation and development (*Nkx1-2* and *Satb2*) were unmethylated, suggesting that these two subtypes represent mature enterocytes and less differentiated cells, respectively (Fig. 2d). Within the immune group, we inferred that the two groups represented T cells (hypomethylated at promoters of genes including *Itk* and *Cd3* complex members) and B cells (hypomethylated at promoters of genes including *Blnk*) (Fig. 2e). Unsupervised dimension reduction approaches, commonly used in single-cell analyses, can reveal otherwise hidden groups of cells. We used NMF for this task (instead of UMAP or PCA) due to its ability to natively accommodate sparse data. NMF of variably methylated sites perfectly recapitulated the cell type groupings inferred by analysis of cell-type-specific hypomethylated regions and was additionally able to partition doublets and putative G2-phase cells (Fig. 2f, g).

**Figure 2.**
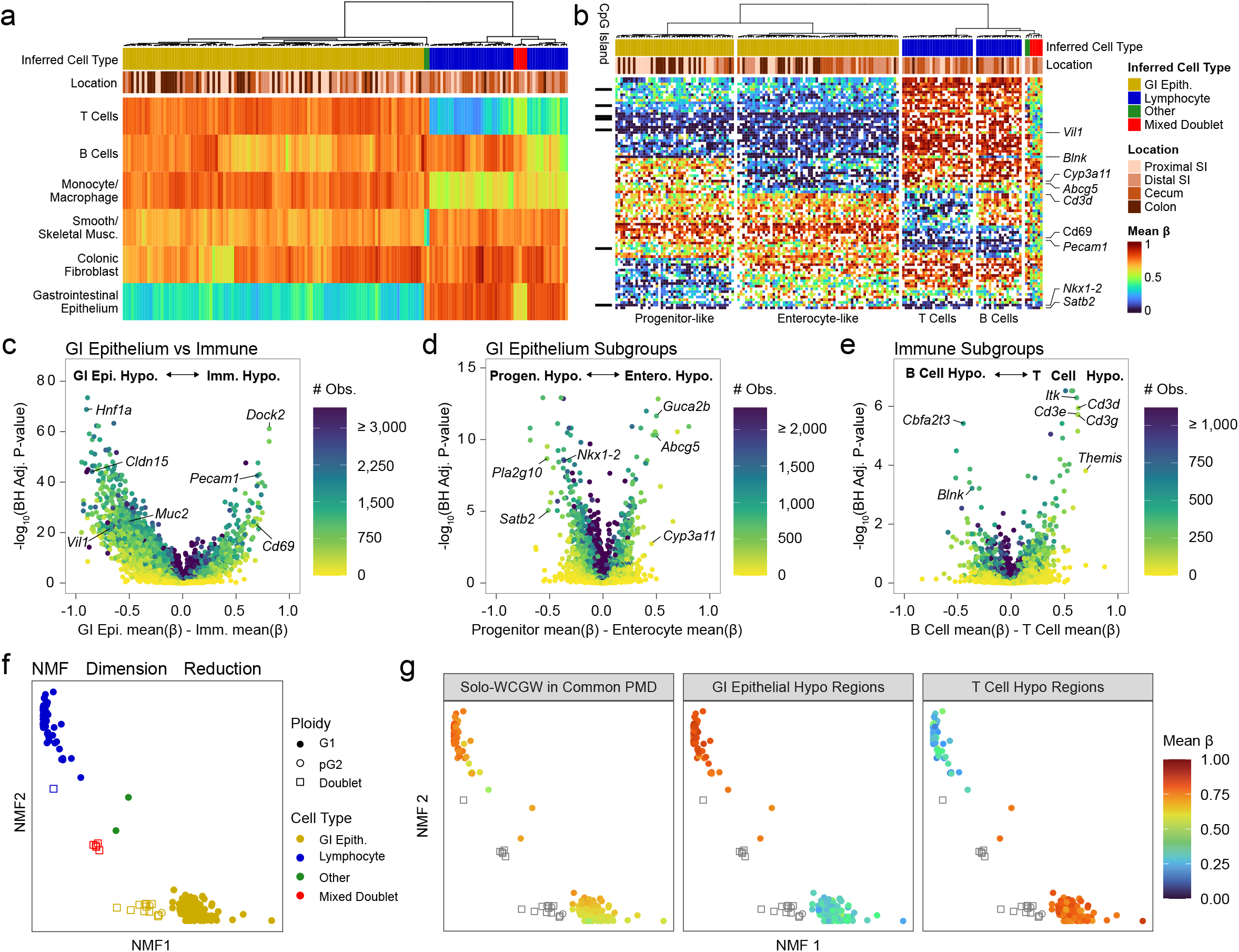
scDEEP-mC captures high-resolution cell type and state information. (a) Unsupervised hierarchical clustering of n=164 high-coverage cells and doublets from primary mouse intestinal epithelium by mean methylation state in cell-type-specific hypomethylated regions^11^. Gastrointestinal epithelial and lymphocytic cell types can be clearly distinguished, as well as several doublets incorporating different cell types with intermediate beta values. (b) Unsupervised hierarchical clustering of differentially methylated promoters exactly recapitulates the cell types found in (a), but with additional granularity and biological insight. Most cell-type-specific promoters do not incorporate CpG islands. (c-e) Volcano plots illustrating differentially methylated promoters between cell groups (P values from unpaired t-tests, subject to Benjamini-Hochberg correction). The number of methylation calls contributing to each test is shown on the color scale. (f) Rank-2 NMF dimension reduction of raw beta values at 4.7 million variably methylated CpGs in n=175 cells perfectly recapitulates the cell types discovered in (a). Note that doublets and putative G2-phase cells (see Fig. 4c) are segregated from their respective cell types. (g) Cell-type-specific hypomethylation and solo-WCGW methylation appear to contribute to NMF clustering.

scDEEP-mC is especially well suited to investigate complex and poorly characterized DNA methylation phenomena such as hemi-methylation. scDEEP-mC profiles hemi-methylation directly, without requiring cellular perturbations^12^, specialized library construction techniques^13,14^ or indirect inference^8^. By examining heterozygous single-nucleotide polymorphisms (SNPs), methylation calls can be assigned to alleles, allowing quantification of hemi-methylation (asymmetric methylation on the reference and complement strands of the same allele) and investigation of allele-specific methylation phenomena at the single-cell level. To add allele resolution to our DNA methylation data, we developed a bisulfite-aware, fully-reproducible Nextflow15 pipeline (see Methods), which (in contrast to existing algorithms^16^) does not require a database of known heterozygous SNPs, creation of a custom reference, or time-consuming re-alignment (Fig 3a).

**Figure 3.**
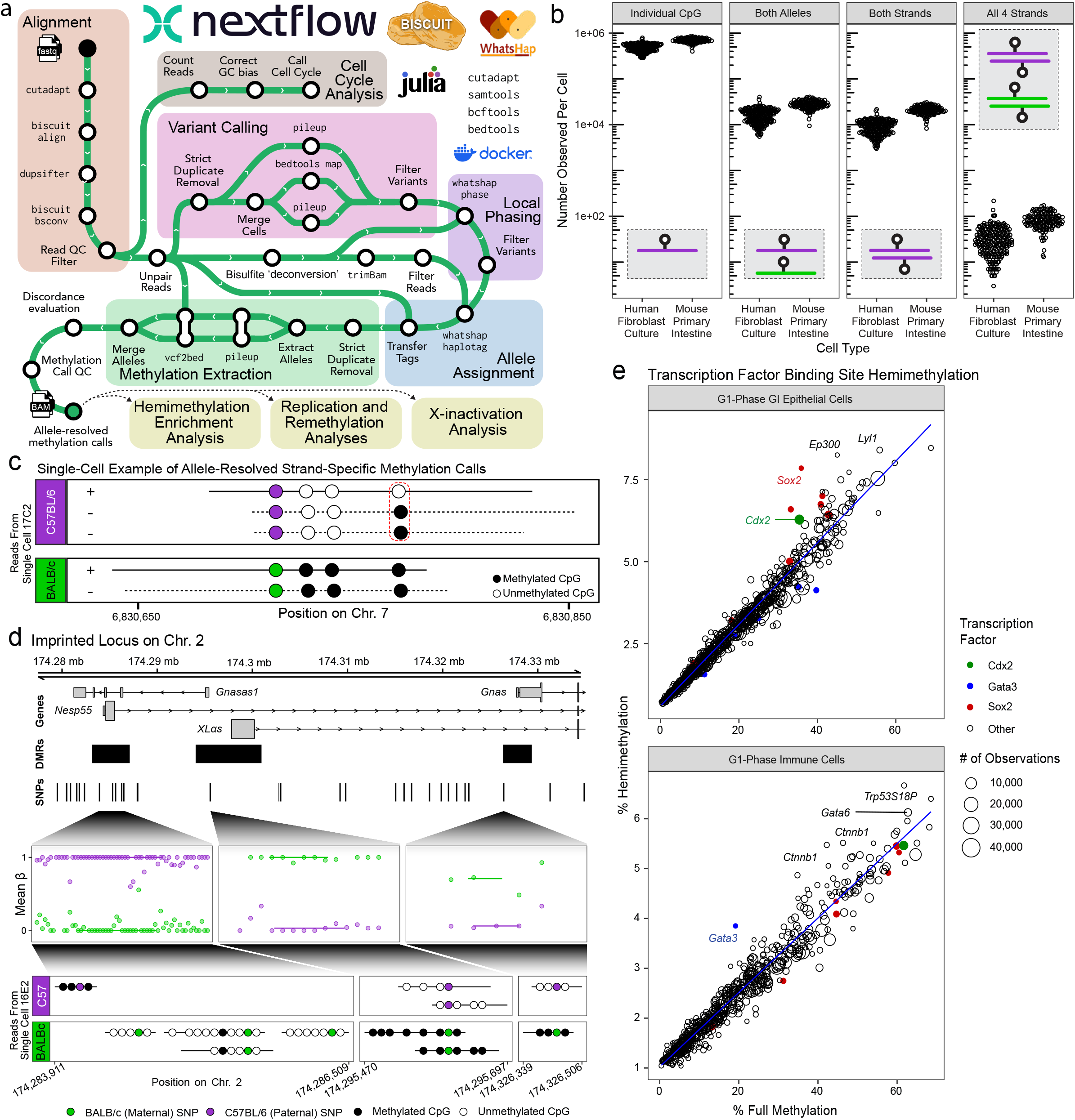
scDEEP-mC facilitates high-resolution, genome-wide analysis of allele-resolved methylation, including hemi-methylation. (a) Overview of allele-resolved methylation analysis. High-quality heterozygous SNPs are discovered and used to assign reads to local alleles (see Methods). (b) Allele-resolved CpG coverage. In primary mouse cells (known SNPs) and human cells (SNPs discovered *de novo*), approximately 1 million CpGs are assigned to an allele. Approximately 10,000 loci per cell have two-strand coverage, allowing for hemi-methylation measurement. (c) Example locus showing reads from a single cell providing information from all 4 strands (reference and complement, each allele), highlighting hemi-methylation and allele-specific methylation. SNPs are shown relative to the reference sequence. (d) Example imprinted (Gnas) locus. Reads from one cell define all three differentially methylated regions. (e) Enrichment of hemi-methylation in TFBS. Hemi-methylation content is plotted against (symmetric) methylation frequency, since the absolute quantity of hemi-methylation is strongly correlated to TFBS methylation level.

We tested our allele-resolved methylation (ARM) analysis on primary intestinal epithelial cells from an F1 cross between two inbred mouse strains (C57BL/6 and BALB/c), as well as cultured primary human fibroblasts. In both groups, we generated allele-resolved methylation calls for nearly 1 million CpGs per cell, including approximately 10,000 symmetrically covered CpGs (permitting analysis of hemi-methylation [Fig. 3b, c]). ARM analyses highlight the high coverage of scDEEP-mC; for example, a single cell is sufficient to define all three differentially methylated regions (DMRs) on both alleles of the imprinted Gnas locus (Fig. 3d).

Since scDEEP-mC can measure two-strand methylation states in single cells, it enables *post hoc* population-specific analyses of hemi-methylation (Fig. 3e). We measured overall methylation and hemi-methylation at TFBS in each of the major cell groups described in Fig. 2b and found that Cdx2 binding sites were more often hemi-methylated than expected in gastrointestinal epithelial cells, while Gata3 binding sites were highly hemi-methylated in immune cells. Additionally, Sox2 binding sites had relatively high rates of hemi-methylation in gastrointestinal epithelial cells. This finding is particularly interesting considering that the ‘super-pioneer’ demethylating activity of Sox2 has been demonstrated to be mediated through replication and passive dilution of methylation, rather than active demethylation by TET enzymes^17^.

Previous work has shown that CpGs in late-replicating loci are prone to DNA methylation loss over successive mitotic divisions^18,19^ and suggested that this may be due to incomplete remethylation after semi-conservative replication of 5-methylcytosine (Fig. 4b)^20^. We measured the relative amounts of unmethylation, hemi-methylation, and full methylation in 13 genomic regions delineated by replication timing^21^ in low and high-passage (TERT-immortalized) human fibroblasts (Fig. 4a). This demonstrated that later-replicating loci have higher rates of hemi-methylation in early passage cells. In late-passage cells, these late-replicating loci have lower absolute levels of methylation, but the relative amount of hemi-methylation does not change appreciably, suggesting that these loci are indeed less amenable to re-methylation after replication.

**Figure 4.**
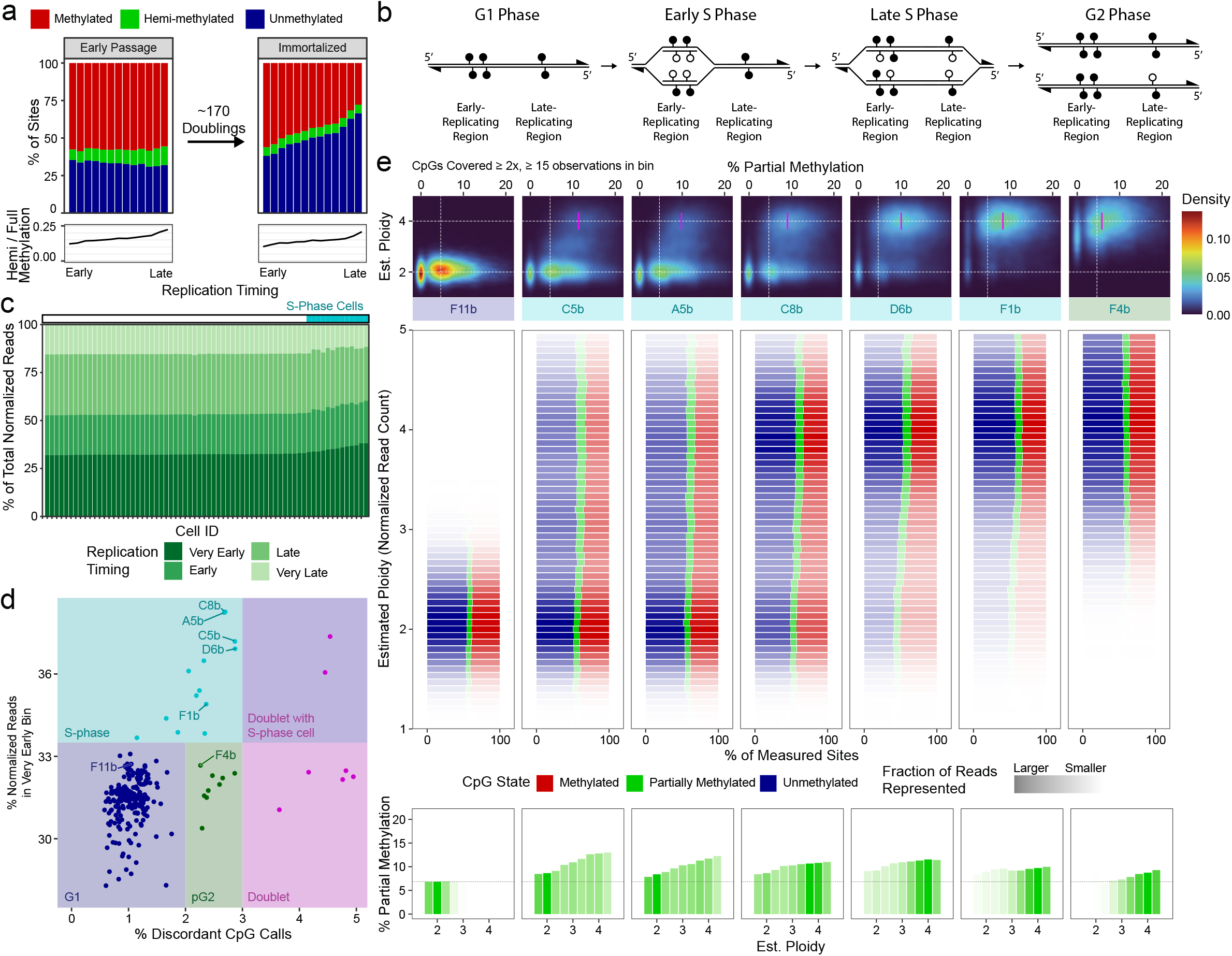
scDEEP-mC enables profiling of replication-induced partial methylation and re-methylation in actively replicating cells. (a) Quantification of two-strand methylation state across the replication timing spectrum in human fibroblasts. In early-passage cells, more hemi-methylation is present in late-replicating regions than in early-replicating regions. Over many mitoses, incomplete maintenance of methylation at these loci leads to loss of methylation. Notably, the amount of hemi-methylation relative to symmetric methylation remains relatively unchanged. (b) Diagram illustrating methylation dynamics during replication. Semi-conservative replication of 5-methylcytosine results in hemimethylation, which is slowly resolved by maintenance methylation. (c) Identification of replicating cells. The fraction of normalized reads (see Methods) assigned to very early, early, late, and very-late-replicating regions of the genome is shown for a selection of cells, sorted by the fraction of reads in the ‘very early’ bin. S-phase cells have more reads in earlier-replicating bins. (d) The fraction of reads in very-early-replicating regions (Fig. 4b) is plotted against the fraction discordant methylation calls for each cell. Cells in S-phase have more reads in earlier-replicating regions of the genome and higher discordance; putative G2-phase cells (pG2) exhibit higher discordance, but even read distribution. Doublets exhibit very high discordance. (e) Distribution of methylation states in replicating cells. Seven single cells in various stages of replication are shown, ordered by their replication progress. For each cell, the estimated ploidy and proportion of CpGs with partial methylation was calculated for 50kb windows across the genome. Top panels show distribution of bins by estimated ploidy and fraction partial methylation; the modal % partial methylation for replicated regions is marked in magenta. Middle panels show the distribution of methylation states for each cell, aggregated by estimated ploidy; the distribution of reads is depicted by the opacity of the bar (regions with more reads are more opaque). Bottom: same data as middle panels, focusing on % partial methylation.

To further investigate the relationship between replication and DNA methylation, we sought to identify actively replicating cells. We reasoned that cells in S-phase would have more reads aligned to early-replicating regions of the genome (Fig. 4c), as well as higher methylation discordance due to incomplete remethylation. We successfully identified cells which met these criteria, as well as putative G2 cells with higher methylation discordance but an even distribution of reads across the genome (Fig. 4d). We then quantified the methylation discordance and normalized read count in 50kb bins across the genome in cells in various stages of replication (Fig. 4e). As expected, newly replicated bins have higher discordance than unreplicated bins due to incomplete remethylation of the daughter strand. As cells progress through S-phase and into G2 phase, this discordance decreases (Fig. 4e).

Existing methods for studying X-inactivation in single cells rely on transcriptomics^22^, which generates no data from the inactive chromosome. In contrast, scDEEP-mC is uniquely able to characterize the epigenetic state of both X alleles in single female cells, allowing us to resolve mechanisms of X-inactivation imbalance that are inscrutable to RNA-based analysis. We utilized cells from a C57BL/6xBALB/c mouse to phase SNPs on chromosome X to the maternal or paternal allele. This allowed us to determine which X was inactive in a particular cell (Fig. 5a). CpG islands (CGIs) and promoters were generally methylated on the inferred inactive X chromosome (Xi) and unmethylated on the inferred active X chromosome (Xa). Some CGIs were methylated on both alleles, while only the CGIs proximal to the Xist and Dxz4 macrosatellite loci^23^ were methylated on Xa and unmethylated on Xi (Fig. 5b). Two regions were identified that were unmethylated on both alleles; of these, *Nap1l3* is known to escape X-inactivation^24^, while the *Pcdh11* CpG island is in a gene body. Likewise, the promoters of inactivated genes on Xi exhibited higher methylation levels than those on Xa, while gene bodies of these genes displayed slightly higher methylation levels on Xa (Fig. 5c).

**Figure 5.**
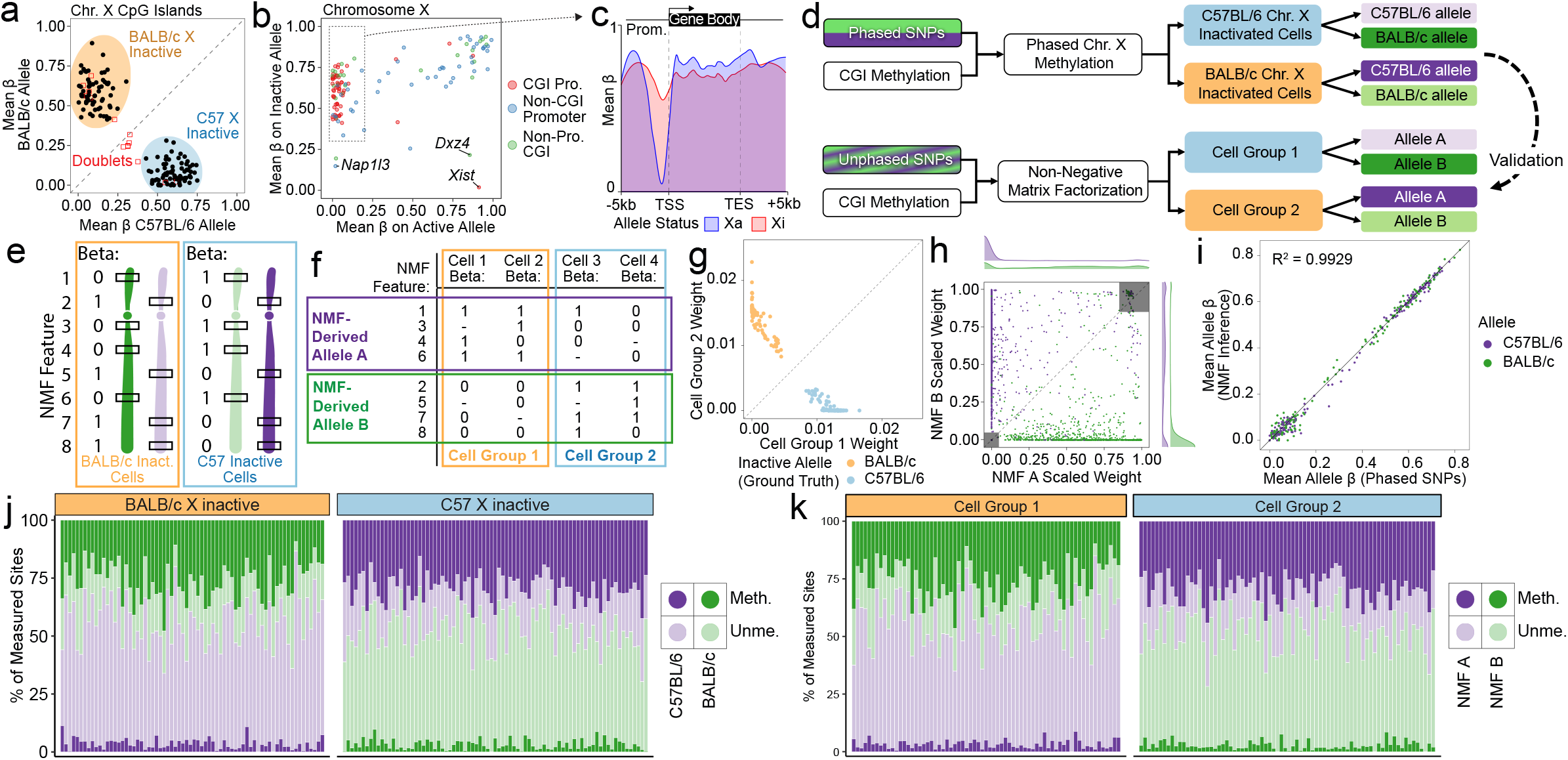
scDEEP-mC enables quantification of whole-chromosome X-inactivation epigenetics in single cells, with or without phased SNPs. (a) Mean CpG island methylation on the active and inactive X chromosomes in n=151 cells from primary mouse intestine. (b) Mean methylation on the Xa and Xi (as defined in (a)) for promoters and CpG islands on chromosome X, across all single cells. (c) Metagene plot showing methylation levels across genes subject to X-inactivation (outlined in (b)). (d-f) NMF can be used to model X-inactivation in populations of cells by bipartitioning highly correlated methylation states across cells and chromosomes. High confidence phased SNPs from the Mouse Genome Project were used as the ground truth. (g, h) NMF analysis almost perfectly recapitulates ground-truth data, correctly assigning cell X-inactivation state and allele membership. (i) Mean methylation values of loci on methylation-derived alleles computed by NMF correlate very well with the corresponding ground-truth values. (j, k) CpG island methylation distribution is shown for each cell; colors correspond to allele, while saturation denotes methylation state. Ground truth data from phased SNPs (j) or NMF-derived alleles (k) are shown.

Since the behavior of CGI methylation across Xi is highly correlated, we investigated whether we could infer X-inactivation status analytically (Fig. 5d-f), avoiding the necessity of phasing SNPs across the entire chromosome (which is infeasible with short-read sequence data). We used rank-2 NMF of raw allele-resolved chromosome X beta values to bipartition the dataset and found that the resulting groups of cells and alleles almost perfectly recapitulated the ground truth dataset (defined by strain-specific SNPs) (Fig. 5g-k). This advance opens the opportunity to analyze X inactivation states in any heterogeneous population of female cells, even when SNP phasing is not available.

We applied our analytical approach to cultured female human fibroblasts to examine the effect of extended culture on X inactivation (Fig. 6a). When we measured the standard deviation of allele-resolved beta values in CGIs across cells, we noticed that early-passage cells exhibited high variance on chromosome X, reflecting the random assortment of X-inactivation states within the population. The late and (TERT-immortalized) very-late passage cells, however, had diminished allele-resolved methylation variance on chromosome X, indicative of a loss of CGI methylation diversity between cells (Fig. 6b). Several mechanisms could be responsible for this phenomenon, including population-level loss of CGI methylation on Xi, population-level chromosomal loss of Xi (with or without duplication of Xa), and evolutionary drift in the population favoring one X-inactivation state (Fig. 6c). Notably, both loss of Xi and evolutionary drift result in a homogenous population of cells expressing transcripts from one copy of one parent’s X, making these scenarios impossible to distinguish with RNA-based single-cell X-inactivation analyses. To distinguish between these scenarios, we used NMF to identify two highly correlated groups of cells and loci on chromosome X in early-passage cells (which we inferred to represent X-inactivation states). We then measured the number of observed loci and methylation state of each of these NMF-derived alleles in late and very-late passage cells.

**Figure 6.**
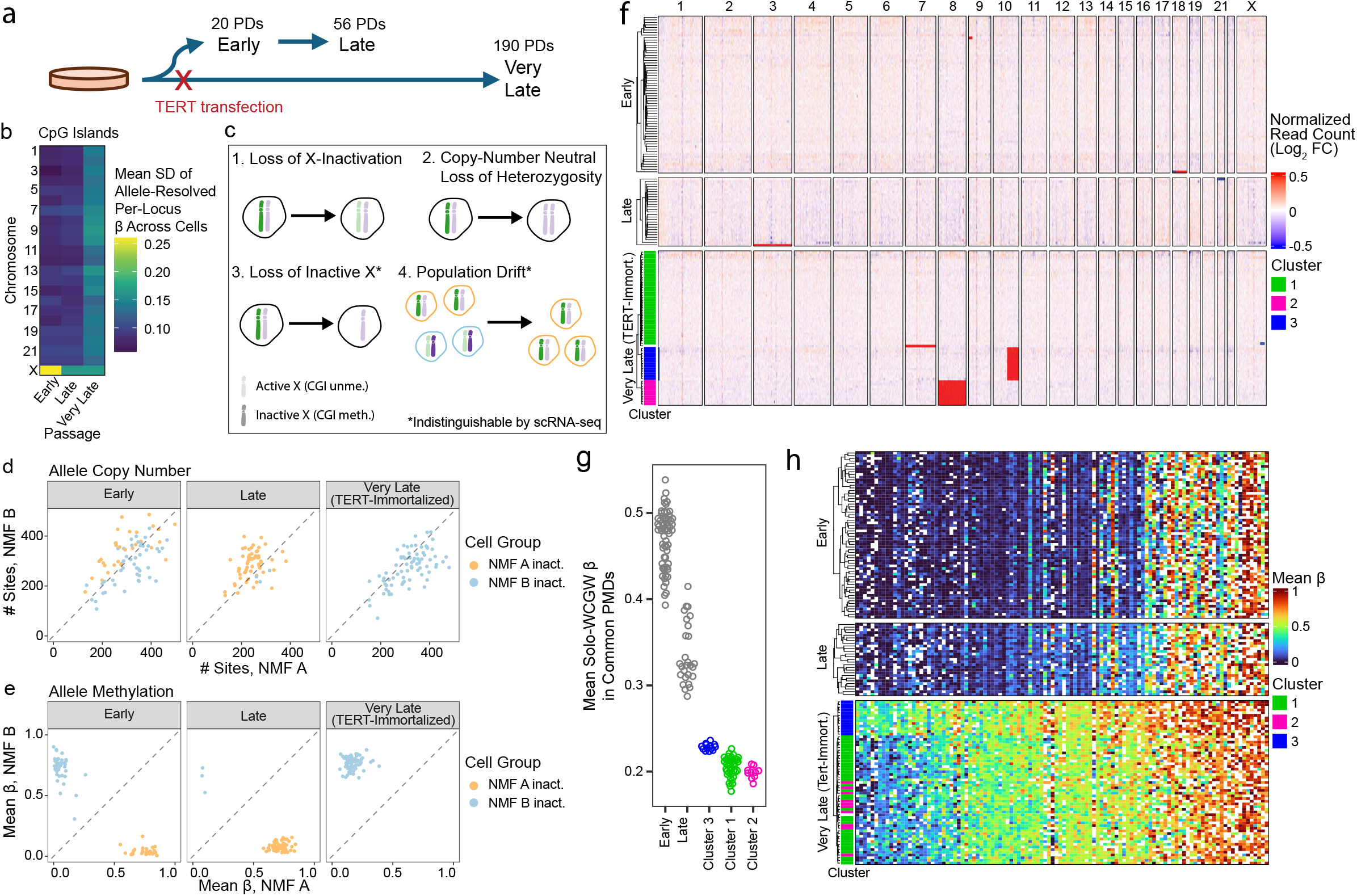
scDEEP-mC illuminates X-inactivation changes and population heterogeneity in cultured human fibroblasts. (a) Experimental overview. AG06561 cells were obtained from Coriell; a subculture was immortalized via TERT transfection and cultured for ∼1 year. A parallel culture was passaged until senescent. (b) High between-cell variability of CGI methylation states on a single allele is noted on chromosome X in early passage cells, indicative of a mixture of X-inactivation states in the population. (c) Loss of variability in CGI methylation on the X chromosome in late and very late-passage populations suggests several possible explanations, some of which are impossible to distinguish via expression-based methods. (d, e) NMF analysis of chromosome X methylation in early-passage cells groups loci into NMF-derived alleles. Analysis of the number of methylation calls for (d) and methylation state on (e) each NMF allele at different passage levels demonstrates that the loss of variability in chromosome X CpG islands at late passage is due to population drift rather than copy number changes or severe X-inactivation dysregulation. (f) Copy number analysis of early, late, and very-late passage cells confirms that no substantive copy number alterations are present on chromosome X and identifies three major subclones in very-late-passage cells (one diploid and two hypodiploid). These clusters have different mean solo-WCGW methylation values (g) as well as differential CpG island methylation state (h), illustrating the power of scDEEP-mC to uncover variation even within relatively homogenous cell populations.

We observed similar numbers of CpG methylation calls from each NMF allele in all passages, indicating that both NMF alleles are present in all passages (Fig. 6d). However, we observed a preferential shift toward methylation of a single NMF allele in late and very-late passage cells, suggesting population drifts in X-inactivation state (Fig. 6e). Read-count-based copy number analysis also demonstrated the absence of substantive copy number alterations on chromosome X in any passage (Fig. 6f). However, we noticed two aneuploid subclones (with chr8 [Cluster 2] and chr10q [Cluster 3] deletions, respectively) in the very-late-passage cells, which exhibited differing levels of solo-WCGW methylation (suggestive of differential replication history [Fig. 6g]) and slight differences in CGI methylation (Fig. 6h), illustrating the power of scDEEP-mC to characterize subtle but important differences in relatively homogenous cell populations.

While many single-cell DNA methylation technologies focus on scaling to very high throughputs^3,25,26^, high-coverage, efficient library generation techniques have the potential to reveal previously hidden insights when coupled with mechanistically-informed analyses. As examples, we demonstrate how scDEEP-mC can offer insight into hemi-methylation, the interaction between replication timing and re-methylation, and whole-chromosome readouts of X-inactivation in primary cells without requiring external manipulations or specialized techniques. scDEEP-mC library construction is straightforward and amenable to combination with other genomic methods, such as STORM-seq^27^, offering the potential for further insights into the relationship between DNA methylation and the transcriptome.

## Acknowledgements

This work was supported by NIH grants R01CA234125 and R01AG084743 awarded to PWL, and R37CA230748 awarded to HS. We thank Zach DeBruine for insightful discussions. We are grateful to Liang Kang and Emily Jung for assistance with animal husbandry and cell culture experiments. We thank the Vivarium (RRID:SCR_023211), Flow Cytometry (RRID:SCR_022685) and Genomics Core (RRID:SCR_022913) resources at Van Andel Institute for their contributions of technical expertise and insights. Computation for the work described in this paper was supported by the High Performance Cluster and Cloud Computing (HPC3) Resource directed by Zach Ramjan at the Van Andel Research Institute.

## Methods

### Primary Cell Culture and Transduction

Primary human fibroblasts (AG06561) obtained from the NIA Aging Cell Culture Repository at the Coriell Institute for Medical Research were maintained in Eagle’s MEM with Earle’s salts, non-essential amino acids (Gibco 11140-050), and 15% v/v fetal bovine serum. Cells were maintained at 37°C, 5% CO_2_, and 21% O_2_. Low-passage primary fibroblasts were transduced with purified lentiviral particles containing expression vectors encoding human telomerase reverse transcriptase as previously described^19^.

Prior to flow sorting, cells were washed twice in PBS and incubated with 0.25% trypsin for 5 minutes, allowing the cells to dissociate from the dish. Growth media was then added to the cells, which were centrifuged at 300 g for 5 minutes. The resulting pellet was resuspended in flow buffer (HBSS without magnesium or calcium, with 5% FBS, 5 mM EDTA, and 1 μg/mL DAPI) with a volume sufficient to achieve 10^6^ cells/mL.

### Intestinal Cell Isolation

Tissue samples were collected from the small intestine, cecum, and colon of 8-week-old CB6F1/J female mice purchased from Jackson Labs (Strain # 100007). For the small intestine and colon, the dissection involved collecting approximately 2 cm sections from the proximal and distal small intestine, and about 4 cm of the mid-colon. The tissue pieces were washed three times with cold PBS, then cut into 2 mm pieces and placed in 15 mL of 5 mM EDTA/PBS with 22.5 µL of 1 M DTT. The tissue was triturated, transferred to 15 mL of fresh 5 mM EDTA/PBS with 22.5 µL of 1 M DTT and incubated for 15 minutes. The tissue was then washed with cold PBS twice to wash out EDTA. The tissue was resuspended in 10 mL collagenase IV solution (Stemcell Technologies 07909) with 100 µL DNase I (Ambion AM2235) and 10 µL ROCK inhibitor Y-27632 (Sigma-Aldrich Y0503), then incubated for 10 minutes at 37 °C. The intestine was then triturated four times with a total of 40 mL cold PBS, and the filtrate was collected by filtering the supernatant through 100 µm, 70 µm, and 40 µm filters. The supernatant was spun at 300g for 5 minutes at 4 °C. The pellet was then resuspended in 1 mL flow buffer (10 mL of PBS with magnesium and calcium, 200 µL FBS, 100 µL DNase I, and 10 µL Y-27632); cells were then counted using a Countess II instrument (Thermo Fisher). 10^6^ cells were transferred into 100 µL of flow buffer, and Fc receptors were blocked with 0.5 µg of TruStain FcX PLUS (anti-mouse CD16/32, BioLegend Cat. No. 156603) for 10 minutes on ice. Without washing the cells, 0.5 µg PE anti-mouse CD45.2 recombinant antibody (BioLegend Cat. No. 111103) and 1 µg Alexa Fluor 647 anti-mouse CD326 (EpCAM) antibody (BioLegend Cat. No. 118211) were added. Cells were incubated in the dark, on ice, for 30 minutes. Cells were then washed twice (2 mL flow buffer, centrifuge at 300g for 5 minutes at 4 °C). Finally, cells were resuspended in 500 µL flow buffer; DAPI (1 µg/mL) was added prior to flow sorting.

### Cell Sorting

Flow sorting was performed using a BD FACSymphony™ S6 Cell Sorter. Live cells were selected for sorting based on scatter properties and DAPI negativity. BD FACSDiva™ Software and BD StepSort were used for data acquisition and 96-well sorting. Flow plots were generated using FlowJo v10.9.

### scWGBS Library Preparation, Quality Control, and Sequencing

Cells were FACS sorted into 3µL of freshly prepared CT Conversion Reagent (Zymo EZ DNA Methylation-Gold Kit) in semi-skirted low-DNA-binding 96-well plates (Eppendorf twin.tec LoBind). Immediately after sorting, lysis and bisulfite conversion was performed by heating to 98°C for 8 minutes, followed by three conversion steps (64°C for 1 hour) alternating with denaturation steps (98°C for 2 minutes). 1µL of 10M sodium hydroxide was added to facilitate desulfonation; the reaction was then incubated at 37°C for 15 minutes. 96 µL of 10.4 mM Tris-HCl was then added to neutralize the reaction mixture and lower the concentration of sodium bisulfite, for a final volume of 100 µL.

Primer composition was determined by counting the proportion of 2-base kmers in the genome, then changing all Cs to Ts except in the CpG context. This yielded an estimate of approximately 49% T, 30% A, 20% G, and 1% C (in the CpG context) in the converted genome. Thus, our first-strand random primer is composed of 50% A, 30% T, and 20% C, plus a 1% spike-in of G exclusively in CpG context; the second-strand random primer is the complement of this composition.

The first strand primer mix was prepared, consisting of 0.5 µM Oligo 1 and 0.625 nM each G-poor CpG spike-in primer. Next, 14 µL of first strand random priming mixture (170 µL first strand primer mix, 170 µL 10 mM dNTPs, 1166 µL 10x Blue Buffer) was added to the reaction, which was incubated at 65°C for 3 minutes, then chilled to 15°C. 1 µL (50U) of high-concentration klenow 3’-5’ exo-polymerase (Qiagen) was added, and a linear amplification cycle was initiated (15°C for 5 minutes, slow heating [0.1°C/sec] to 37°C, hold at 37°C for 5 minutes, chill to 15°C for 10 seconds, heat to 95°C for 1 minute). The reaction was then chilled to 15°C, and 2.5 µL of master mix (740 µL first strand primer mix, 74 µL 10 mM dNTPs, 185 µL 10x Blue Buffer, 370 µL high-concentration klenow exo-, 481 µL water) was added before another linear amplification cycle was performed, using the same incubation steps as above; this process was repeated for a total of 7 linear amplification cycles. For the last cycle, 5 µL of master mix was added instead of 2.5 µL. Finally, the reaction was chilled to 15°C for 5 minutes, slowly heated (0.1°C/sec) to 37°C, held at 37°C for 90 minutes, and chilled to 4°C. Prior to second strand synthesis, 2 µL (40 U) Exo I (NEB) was added, and the reaction was incubated at 37°C for 1 hour to digest single-stranded DNA fragments. A bead cleanup (Beckman Coulter SPRIselect) was then performed using 106 µL of beads per reaction (0.8x ratio), eluting into 40 µL of water.

To incorporate the second sequencing primer, a second strand primer mix was prepared, consisting of 10 µM Oligo 2 and 12.5 nM each C-poor CpG spike-in primer. 9 µL of second-strand master mix (206 µL 10 mM dNTP, 206 µL second strand primer mix, 515 µL 10x Blue buffer) was then added, and the reaction was incubated at 95°C for 1 minute, then cooled to 4°C. Next, 2 µL (100U) of klenow exo-was added, and the reaction was held at 4°C for 5 minutes before being slowly heated (0.1°C/sec) to 37°C, then held at 37°C for 90 minutes. The reaction was cooled to 4°C and a second bead cleanup was performed, using 40 µL of beads (0.8x ratio) and eluting into 23 µL of water. 23 µL of KAPA HotStart ReadyMix master mix (Roche) and 2 µL of xGen UDI indexing primers (IDT) were added, followed by PCR amplification (95°C for 2 minutes; 12-15 cycles of melting [94°C for 80 seconds], annealing [62°C for 30 seconds], and extension [72°C 40 seconds]; 72°C for 3 minutes). A final bead cleanup was performed using 35µL of beads (0.7x) and eluting into 21 µL of storage buffer (10 mM Tris, 0.1 mM EDTA, pH 8.0).

After preparation, the concentration and fragement size distribution of each library was quantified using the Quant-iT PicoGreen dsDNA kit (Invitrogen) and High Sensitivity D1000 ScreenTape Assay (Agilent), respectively. Libraries (typically 30-35 per pool) were pooled in an equimolar fashion and the pool was purified with a 0.8x ratio bead cleanup. Libraries were sequenced to a depth of approximately 30M 150bp paired-end reads using an Illumina NovaSeq 6000, with 10% PhiX spike-in.

### Sequence Data Processing

Base calls were demultiplexed using `bcl2fastq`, allowing for one mismatch, by the sequencing provider. Adapters were trimmed using `cutadapt`^28^ with the parameters `--overlap 1 -a AGATCGGAAGAGC -A AGATCGGAAGAGC --error-rate 0.1 --trim-n --minimum-length 20 -- nextseq-trim 20 -n 2`. Paired, trimmed reads were aligned using `Biscuit`^10^ in a paired directional fashion (`biscuit align -b1 /path/to/ref read_2 read_1`). Note that read 2 is supplied before read 1 to allow for the correct directionality in alignment. Singleton reads (in which one read of a mate pair was discarded by `cutadapt`) were aligned in a single-ended, directional fashion (`biscuit align - b 3 /path/to/ref read_1` or `biscuit align -b 1 /path/to/ref read_2`). For each cell, all three alignments (paired-end, single-end read 1, single-end read 2) were merged and then duplicate marked using `dupsifter`^29^ in single-end mode. The resulting alignments are referred to as ‘raw’ alignments.

Raw alignments were subsequently cleaned by removing reads with more than 1 CpY retention event (using `biscuit bsconv`), reads with MAPQ < 40, and unmapped, secondary, duplicate, or vendor-QC-failed reads (`samtools view -q 40 -F 0×704`)^30^. Regions of anomalously high coverage were then identified (as below) and removed to produce ‘polished’ alignments.

### Identification of Anomalous High-Coverage Regions

The genome was divided into 100bp windows, tiled every 50bp across the genome. For each cell, the number of reads overlapping each bin by at least 50% was tallied. Regions were flagged as anomalous if the number of reads in that bin was above the 80^th^ percentile for that cell; anomalous regions were extended until the number of reads in the bin dropped below the 67^th^ percentile for that cell. Flagged regions ≤ 250bp apart were then merged. Anomalous regions were identified as bins that were flagged in > 90% of cells. Finally, anomalous regions < 250 bp apart were joined together to create a list of regions to exclude.

### Methylation calling

Methylation data was extracted from ‘polished’ alignments using `biscuit pileup` with the flags `-5 10 -3 1 -p`, then summarized using `biscuit vcf2bed -t CG -k 1` and `biscuit mergecg CpN`. retention was calculated using `biscuit qc`. Subsequent analyses were performed using `R` (version 4) as detailed below.

### Quality Control

Quality evaluation was performed using `FastQC`^31^ (on raw and trimmed FASTQ files) and `biscuit qc` (on raw and polished alignments). `MultiQC`^32^ was used to summarize QC results. Low-efficiency cells were excluded if they covered fewer than 20% of CpGs or were sequenced to > 55M reads (human) or > 80M reads (mouse).

### Comparative Analysis

Publicly available data was downloaded from the SRA and NGDC. `fastqc` and `multiqc` were used to assess adapter content. Adapters were trimmed using `cutadapt` with the parameters `--error-rate 0.1 --trim-n --minimum-length 20 -n 2 --overlap 1`, either `--quality-cutoff 20` or `--nextseq-trim 20` (if the data was generated on a sequencer using 2-color chemistry), and additional parameters as described in the table below:

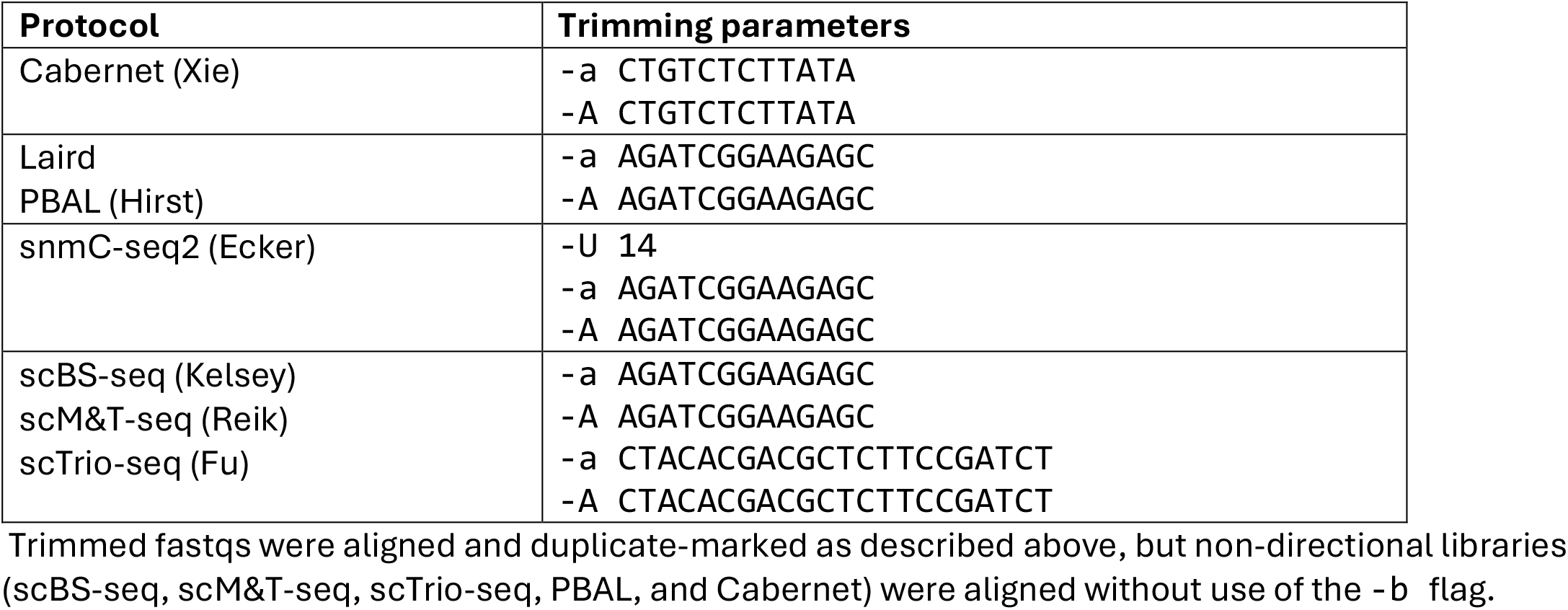

| Protocol | Directional? | Read 1 mapped to: |
| --- | --- | --- |
| Laird<br>snmC-seq2 (Ecker) | Yes | Daughter strand |
| scBS-seq (Kelsey)<br>scM&T-seq (Reik)<br>scTrio-seq (Fu)<br>PBAL (Hirst)<br>Cabernet (Xie) | No | N/A |

Per-protocol identification of anomalous high-coverage regions was performed as described above. Subsequently, aligned reads were filtered as described above, with no protocol-specific changes.

Methylation calling was performed on ‘polished’ alignments using `biscuit pileup` with the `-p` flag set. The following parameters were set on a per-protocol basis to mitigate the effects of random-priming-induced bias in methylation calls:

| Protocol | 5' bases ignored | 3' bases ignored |
| --- | --- | --- |
| Laird | 9 | 0 |
| snmC-seq2 (mouse) | 10 | 6 |
| snmC-seq2 (human) | 9 | 3 |
| scM&T-seq | 6 | 6 |
| scTrio-seq | 9 | 1 |
| scBS-seq | 9 | 6 |
| PBAL | 6 | 1 |
| Cabernet | 9 | 1 |

### Methylation Concordance

Methylation concordance was calculated as follows. For each cell, CpG sites covered more than once were classified as concordant (β exactly 0 or 1) or discordant (0 < β < 1). Then, the ‘discordance metric’ was calculated as:

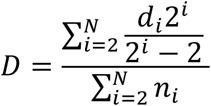

Where *D* is discordance, *d*_*i*_ is the number of discordant sites at coverage depth *i*, and *n*_*i*_ is the total number of sites covered *i* times. Intuitively, this can be conceptualized as calculating the proportion of discordant sites at a given coverage depth, normalized by the probability of observing discordance given random assortment of methylation calls; then taking the mean across all observed depths, weighting by the number of observations at that depth. *D* may be calculated for either allele-resolved or summarized methylation data. In the former case, *D* represents the accumulation of polymerase, sequencing, and allele assignment errors, while in the latter case, *D* also incorporates hemi-methylation and allele-specific methylation.

### Doublet Identification

Libraries generated from K562 cells and tumor samples were excluded, as we found that their hypotriploid karyotype or copy number alterations, respectively, produced artificially inflated coverage due to an increased amount of input DNA. scDEEP-mC libraries were excluded if their allele-resolved CpG discordance rate was > 2%. Libraries from other protocols were excluded if they were not derived from embryonic stem cells and had a CpG discordance rate > 15%.

### Bisulfite Conversion

Base-resolution CpA, CpC, and CpT retention rates were calculated using `biscuit qc CpY`. retention was calculated as the mean of CpC and CpT retention rates.

### Sequencing Efficiency

Adapter trimming yield was computed by calculating the total number of bases in the trimmed fastq files, divided by the expected number of sequenced bases (reported sequenced reads * read length (* 2 if paired)). For subsequent steps, 2 million (paired-end, if applicable) reads were randomly sampled from each fastq file using `seqtk`^33^. These reads were extracted from the aligned .bam file, and duplicate flags were unset. The number of aligned bases in the subset reads was tallied to calculate the alignment yield. Duplicates were then re-marked (using `dupsifter`, as above) in each subset of reads. Reads with > 1 CpY retention event were then discarded (using `biscuit bsconv`, as above). Next, unmapped, vendor-QC-fail, secondary, and low-mapping-quality alignments were discarded (`samtools view -F 0×304 - q 40`). Next, duplicate reads were discarded. After each filtering step, the number of bases in the remaining filtered alignments was tallied using `samtools depth`. Finally, the number of bases covered by the filtered reads was tallied using `samtools coverage`.

### Cell-Type-Specific Hypomethylated Regions

A table of 50286 cell type-specific unmethylated markers (top 1000 for each cell type) was downloaded from^11^. These regions were lifted over from the hg19 source genome to the mm10 genome using the `liftover` tool and hg19ToMm10.over.chain.gz file provided by UCSC. For each cell, the mean β value of all covered CpGs inside all regions specific to a particular cell type (or group of cell types) was calculated. The mean β values for each cell for the “Blood-T”, “Blood-B”, “Colon-Ep:Gastric-Ep:Small-Int-Ep”, “Skeletal-Musc:Smooth-Musc”, “Blood-Mono+Macro”, and “Colon-Fibro” regions were extracted, and cells and regions were hierarchically clustered based on mean β values using Ward’s algorithm (`method = “ward.D2”`). Data was visualized using the `ComplexHeatmap` package^34^. Cells were classified based on their mean β values for the ‘Blood-T’ and ‘Colon-Ep:Gastric-Ep:Small-Int-Ep’ cell types, based on the following criteria:

| | Gastrointestinal Epithelial<br>mean $\beta > 0.65$ | Gastrointestinal Epithelial<br>mean $\beta < 0.65$ |
| --- | --- | --- |
| 'Blood-T' mean<br>$\beta > 0.65$ | Other | GI Epithelial |
| 'Blood-T' mean<br>$\beta < 0.65$ | Lymphocyte | Doublet of two different<br>cell types |

### Solo-WCGW Methylation

Solo-WCGW CpGs in common partially methylated domains in the mm10 and hg38 genome were downloaded from^18^. The mean β value over all covered CpGs in these regions was calculated for each cell.

### Promoter Methylation

For each gene in the GENCODE VM23 (UCSC knownGene) annotation, the region 1.5kb upstream and 200bp downstream of the start site were taken as the promoter. The mean β value of all covered CpGs in each promoter was calculated for each cell. For visualization, promoters with > 25% of CpGs covered in ≥ 60% of cells, and having a standard deviation of > 0.2 across all cells, were selected. Cells and promoters were hierarchically clustered based on mean β values using Ward’s algorithm. For differential methylation analysis, the cells were classified as ‘immune’ or ‘epithelial’ as described under ‘Cell-Type-Specific Hypomethylated Regions’. An unpaired t-test was used to compare mean β values of each promoter in all cells in each group; p-values were corrected using the Benjamini-Hochberg algorithm. Promoters overlapping CpG islands (as defined by^35^) by at least 200 bp were annotated appropriately.

### Non-negative Matrix Factorization

For each CpG, the number of cells covering that CpG and the standard deviation (SD) of β value across all cells was calculated. CpGs above the 25^th^ percentile for coverage and 67^th^ percentile for SD were selected for NMF analysis. β values for each cell were accumulated into a vector; a complementary ‘unmethylation’ vector (1-β) was appended to this vector to prevent subsequent normalization from being affected by methylation state. β values of 0 were replaced with 1×10^−12^ and missing values were replaced with 0. The vectors were concatenated as the columns of a matrix, which was column-normalized and log-transformed. The resulting matrix was subjected to rank-2 NMF, masking zeros, as implemented by the `RcppML` package^36^.

### Variant Discovery from scWGBS data

Due to the nature of scDEEP-mC (and other methods involving multiple rounds of random priming), it is possible for multiple library molecules to be generated from the same fragment. This results in a situation where both first-in-pair reads have the same start position, but their mates have differing start points. Generally, duplicate marking tools would not flag these pairs as duplicates. However, we found that including such read pairs led to a higher rate of false positive variant calls. Thus, for the purposes of variant calling, we adopted a stricter duplicate marking approach by first ‘unpairing’ all reads (unsetting all mate-related flags and tags, while tagging the read name with the read number), then re-marking duplicates in single-end mode. These strictly-deduplicated reads were merged for pseudo-bulk variant calling or taken separately for single-cell variant calling. All variant calling was performed using `biscuit pileup` with the flags `-5 10 -3 7 -p -n 7`. Regions that harbored an unusual amount of low-mapping-quality reads also yielded many false positive variant calls. To identify these regions, the mean mapping quality of all strictly-deduplicated reads was calculated for non-overlapping 500bp bins, and bins with a mean MAPQ < 45 were removed from further consideration. Variants were selected if they had an allele frequency between 0.25 and 0.75 in pseudo-bulk analysis, as well as a quorum of cells (≥ 5 cells or ≥ 5% of cells, whichever was greater) directly supporting each variant call. As expected, a substantial number of SNPs were at CpG sites. SNPs that create or destroy CpGs could lead to a methylation bias in the results. To mitigate this, we identified SNPs where the alternate allele created a CpG and used `bcftools consensus` to ‘apply’ these alternate alleles to the references, thus creating a new reference genome with all possible CpGs to use for methylation calling.

### Curation of Mouse SNPs

We conducted 30x whole-genome sequencing of tail clips from 4 siblings of the F1 mouse from which we generated scDEEP-mC libraries. Variant calling was performed using the nf-core/sarek pipeline (version 3.4.2-gb5b766d), using `haplotypecaller`, `bcftools`, `freebayes`, and `strelka` to call variants.

Variants were filtered using appropriate cutoffs for each tool (`haplotypecaller`: quality > 100`, bcftools`: qual > 221, `freebayes`: qual > 25, `strelka`: qual > 25 and passing filters). Variants called by at least 3 callers were selected; selected variants found in at least 3 individuals (autosomes) or 2 out of 3 female mice (chromosome X) were selected. Finally, selected variants were filtered to only include those found in the Mouse Genome Project variant catalog for BALB/c.

### Local Variant Phasing

It is possible for reads to contain more than one high-quality heterozygous SNP. Thus, it is important to ensure that SNPs are not randomly assigned to alleles, which could introduce unnecessary conflicts when trying to assign reads to alleles. To address this problem, we used `whatshap`, a read-backed phasing algorithm^37^, to ensure that variants were assigned to alleles in a locally consistent fashion.

However, `whatshap` is not designed with bisulfite sequence data (and its attendant ambiguities) in mind. To overcome this issue, we ‘deconverted’ our reads using a custom Julia script. This script performs the following actions:

- For bases that are marked as insertions relative to the reference, ambiguous bases (T on the C->T converted strand; A on the G->A converted strand) are replaced with N
- For bases that are at a known, high-quality heterozygous SNP, ambiguous bases are replaced with N and informative bases are replaced with the supported base. For example, a C or T is ambiguous if it is on the C->T converted strand at a C/T SNP. However, a T base at an A/C SNP supports the C allele, and is replaced with a C.
- All other bases are replaced with the reference sequence

Unpaired, strictly-deduplicated, ‘deconverted’ reads were then used to conduct local SNP phasing using `whatshap phase` with the parameters `--only-snvs --ignore-read-groups --merge-reads` with the appropriate genome as a reference. The output of `whatshap phase` was further edited such that singleton SNPs (SNPs which could not be phased into larger blocks) were transformed into singleton phase groups containing only that single SNP.

### Assignment of reads to local haplotypes

The 5’ and 3’ ends of reads may have substantial methylation bias due to imperfect priming. Thus, we manually trimmed reads using `trimBam` (parameters `-L 10 -R 7`)^38^ and used `samtools view` to exclude reads with 7 or more mismatches. Filtered, deconverted reads were then assigned to alleles using `whatshap haplotag` using the locally phased, high-quality heterozygous SNPs previously generated. The allele assignments generated by this tool were transferred to the original reads using a custom Julia script.

### Allele-Resolved Methylation Calls

Reads assigned to each allele were extracted using `samtools view`. Methylation was called against an updated reference containing all possible CpGs (including those created by SNPs), as described above. Methylation calling was performed as above, using `biscuit pileup` and `biscuit vcf2bed`. Finally, a custom Python script was used to annotate the output file with the phase sets that contributed to each allele-specific methylation call. Methylation calls were discarded in the following situations: 1) Methylation call at a CpG-creating SNP on the allele where there is no CpG; 2) reads from multiple phase sets contributed to the methylation call; or 3) methylation calls at CpGs where both bases were SNPs. Per-read SNP and methylation call data was visualized using the R packages `Rsamtools`^39^, `Biscuiteer`^40^, and `Bisplottti`^41^.

### Transcription Factor Binding Site Hemi-methylation

The transcription factor binding sites (TFBS) catalogued in the LOLA^42^ Core ‘codex’ and ‘encodeTFBS’ databases for the mm10 genome were used for this analysis. Two-strand sites (CpGs where both the top and bottom strand were covered, and both strands were confidently assigned to the same microhaplotype) were selected and grouped by their methylation state (symmetrically unmethylated, hemi-methylated, or symmetrically unmethylated). Finally, the fraction of sites in each state overlapping each TFBS region set was calculated; TFBS region sets with ≤ 250 methylation or hemi-methylation events were discarded.

### Read Count Analyses

The number of ‘polished’ alignments falling within 50kb non-overlapping bins was tallied using `bedtools`^43^, then normalized by the total number of polished, aligned reads per cell to yield reads per million mapped (RPMM). Per-bin RPMM biases were normalized as follows: Bins with RPMM < 5^th^ or > 95^th^ percentile for their cell, in > 75% of cells, were excluded, as were bins with > 10% missing data. ‘Ideal’ cells with a uniform read distribution were selected by computing the standard deviation (SD) of RPMM in 5Mb non-overlapping bins. Bins with an SD < 2.5 were selected, and the standard deviation of the mean read count over all bins was calculated for each cell. Cells with a whole-genome SD between 1 and 1.75 were flagged as ‘ideal’ cells. A per-bin correction factor was obtained by calculating the median RPMM for each 50kb bin across all ideal cells; RPMM counts across all cells were normalized by these bin-specific corrections.

### Replication Timing

High-resolution repli-seq data was downloaded from^21^. 50kb non-overlapping bins tiling the genome were assigned a replication timing bin from 1-16 by rounding the mean of the top 3 highest-scoring bins, weighted by the score, to the nearest integer. Replication bins 1, 15, and 16 were excluded from further analysis, as they encompassed disproportionately large or small regions of the genome and may have introduced bias. For each replication timing bin, two-strand methylation states (symmetric methylation, hemi-methylation, or symmetric unmethylation) were tallied across all G1-phase cells in each passage (Fig. 4a).

Replication timing bins were further grouped as follows: “very early” included bins 2-4; “early” included bins 5-8; “late” included bins 9-12; and “very late” included bins 13-14. Cells with > 33.5% of normalized reads in the “very early” bin and < 3% allele-resolved CpG discordance were labeled as ‘S-phase’ cells (Fig. 4d).

For Fig. 4e, normalized read count was converted to estimated ploidy by calculating the mode of the normalized read count density. Normalized read counts were divided by this mode, then multiplied by 2 (if the mode was ≤ 1.1) or 4 (if the mode was > 1.1, or the cell was flagged as G2 phase). The genome was partitioned into 50kb bins, and the β values for all CpGs covered ≥2x were discretized into three groups: unmethylated (β == 0), methylated (β == 1), and partially methylated (0 < β < 1). Bins with < 15 methylation measurements were discarded. The remaining bins were grouped by their estimated ploidy and the total proportion of CpGs in each state was calculated. The opacity of each bin was scaled by its weight, which was calculated as

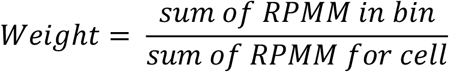

### X-Inactivation Analysis

Per-allele mean β values of CpGs within CpG islands (as defined by^35^) were calculated per cell (Fig. 5a); cells with < 50 methylation calls for each allele were removed. For Fig. 5B, three region sets were considered. *CpG island promoters* were defined as CpG islands within 1.5kb upstream - 1kb downstream of a transcription start site, with the remaining CpG islands being described as *non-promoter CpG islands. Non-CpG island promoters* were defined as regions 1.5kb upstream – 200bp downstream of genes which did not have a CpG island 1.5kb upstream – 1kb downstream of the TSS. All CpGs within a TSS, or within a CpG island/shores (+/-500bp), were averaged across all cells (considering each cell’s X-inactivation state) to produce a mean β for each feature on both the active and inactive X chromosomes. Features with < 100 observations (methylation calls) were discarded.

For Fig. 5C, gene promoters (both CGI and non-CGI) with ≥ 100 methylation calls, a mean β < 0.25 on the active allele, and a mean β > 0.25 on the inactive allele were considered. The positions of CpGs within these genes (and the regions 5kb upstream and downstream) were normalized to align to each other, then binned. Mean β values were calculated for the active and inactive allele within each bin; bins within genes were weighted by the length of the gene (since shorter genes have fewer measurements, and thus more uncertainty). Mean β values were smoothed using a loess algorithm with `span = 0.2` for final visualization.

### NMF of Chromosome X Methylation Data

All allele-resolved methylation calls on chromosome X were considered for NMF. β value complementation, zero-masking, log-normalization, and factorization were performed as described above. After factorization, features > 500bp away from a CpG island were excluded. NMF features having either factor value > 95^th^ percentile for that factor were excluded. Subsequently, factor loadings were scaled to a maximum of 1, and features with a scaled loading < 0.05 or > 0.85 (0.75 for human fibroblasts) in both factors were excluded. Features were assigned to whichever factor had a greater scaled loading. Since each locus is represented by 4 features (one for each allele, one for methylation and one for its complement), CpGs were assigned to NMF-derived alleles by majority vote, discarding CpGs with ties.

### Human Fibroblast Chromosome X Analysis (Fig. 6)

Methylation calls on chromosome X from early-passage cells were subjected to NMF as described above. For each cell, the number of methylation calls assigned to each NMF factor (methylation-derived allele) were tallied to obtain the allele copy number. Likewise, the real β values were summarized to compute allele methylation. Feature assignments derived from NMF analysis of early-passage cells were used to ‘phase’ methylation calls from late- and very late-passage cells.

Normalized read counts were obtained as described above. Cells with < 24% of reads in the ‘very early’ replication timing bin, or < 8M CpGs covered, were discarded (due to high variability in read distribution). Normalized read counts were binned into 0.5 Mb non-overlapping bins and averaged for visualization. Cells were grouped using hierarchical clustering with Ward’s algorithm. The three major clusters in very-late-passage cells were identified and β values in solo-WCGW CpGs in common partially methylated domains^18^ were summarized (as above).

A mean β value was calculated for each CpG island in each cell, discarding CpG islands with ≤ 30% of CpGs covered. CpG islands with a standard deviation > 0.2 across all cells, and < 20% missing data, were selected as ‘variable’. Next, t-tests were computed for each CpG island and major late-passage cluster, comparing its methylation in the cluster to the other two clusters. CpG islands with a Benjamini-Hochberg-adjusted P-value < 0.01 and a mean delta β > 0.15 were selected as ‘differentially methylated’. Cells were hierarchically clustered based on their mean β values at selected CpG islands, and CpG islands were arranged in order of increasing mean β across all very-late-passage cells.

### Primer sequences

All primers were ordered from IDT; the sequences given below are given in IDT ordering format, with the random primer sequence underlined.

#### First strand primers

Oligo1:

ACACTCTTTCCCTACACGACGCTCTTCCGATCT<u>(N1:50200030)(N1)(N1)(N1)(N1)(N1)(N1)(N1)(N1)</u>

G-poor CpG spike-in 1:

ACACTCTTTCCCTACACGACGCTCTTCCGATCT<u>CGHHHHHHH</u>

G-poor CpG spike-in 2:

ACACTCTTTCCCTACACGACGCTCTTCCGATCT<u>HCGHHHHHH</u>

G-poor CpG spike-in 3:

ACACTCTTTCCCTACACGACGCTCTTCCGATCT<u>HHCGHHHHH</u>

G-poor CpG spike-in 4:

ACACTCTTTCCCTACACGACGCTCTTCCGATCT<u>HHHCGHHHH</u>

G-poor CpG spike-in 5:

ACACTCTTTCCCTACACGACGCTCTTCCGATCT<u>HHHHCGHHH</u>

G-poor CpG spike-in 6:

ACACTCTTTCCCTACACGACGCTCTTCCGATCT<u>HHHHHCGHH</u>

G-poor CpG spike-in 7:

ACACTCTTTCCCTACACGACGCTCTTCCGATCT<u>HHHHHHCGH</u>

G-poor CpG spike-in 8:

ACACTCTTTCCCTACACGACGCTCTTCCGATCT<u>HHHHHHHCG</u>

#### Second strand primers

Oligo2: GTGACTGGAGTTCAGACGTGTGCTCTTCCGATCT<u>(N2:30002050)(N2)(N2)(N2)(N2)(N2)(N2)(N2)(N2)</u>

C-poor CpG spike-in 1:

GTGACTGGAGTTCAGACGTGTGCTCTTCCGATCT<u>CGDDDDDDD</u>

C-poor CpG spike-in 2:

GTGACTGGAGTTCAGACGTGTGCTCTTCCGATCT<u>DCGDDDDDD</u>

C-poor CpG spike-in 3:

GTGACTGGAGTTCAGACGTGTGCTCTTCCGATCT<u>DDCGDDDDD</u>

C-poor CpG spike-in 4:

GTGACTGGAGTTCAGACGTGTGCTCTTCCGATCT<u>DDDCGDDDD</u>

C-poor CpG spike-in 5:

GTGACTGGAGTTCAGACGTGTGCTCTTCCGATCT<u>DDDDCGDDD</u>

C-poor CpG spike-in 6:

GTGACTGGAGTTCAGACGTGTGCTCTTCCGATCT<u>DDDDDCGDD</u>

C-poor CpG spike-in 7:

GTGACTGGAGTTCAGACGTGTGCTCTTCCGATCT<u>DDDDDDCGD</u>

C-poor CpG spike-in 8:

GTGACTGGAGTTCAGACGTGTGCTCTTCCGATCT<u>DDDDDDDCG</u>

